# Sodium butyrate inhibits the proliferation of temozolomide-resistant glioblastoma (A172) and lung adenocarcinoma (A549) cells and biphasically regulates EGFR expression

**DOI:** 10.64898/2026.10.26.761832

**Authors:** Minori Ishitsuka, Yuta Hotta, Hiroyoshi Harada, Masayoshi Mori, Yusuke Murata, Kenji Ohe

## Abstract

Glioblastoma carries one of the poorest prognoses of any solid tumour, and once resistance to temozolomide (TMZ) emerges there is little left to offer. Because the TMZ response is governed largely by epigenetic silencing of MGMT, histone deacetylase (HDAC) inhibitors are a rational partner for TMZ. We asked whether sodium butyrate (NaB), a short-chain fatty acid produced by the gut microbiota and a class I/IIa HDAC inhibitor, retains activity against TMZ-resistant cells. In A172 glioblastoma and A549 lung adenocarcinoma cells, NaB changed EGFR expression biphasically: EGFR exon 2–5 mRNA was induced at 0.15 mM in A172 and at 0.30 mM in A549, and was suppressed at higher concentrations, with EGFR protein following the same pattern in A172. NaB reduced proliferation of both lines from 0.31 mM in a concentration-dependent manner, more markedly in A172 than in A549. At concentrations that were inactive alone (NaB 0.30 mM, TMZ 3 µM), the combination inhibited proliferation of A172 on day 3 and of A549 on days 2 and 3. TMZ-resistant sublines (A172 rTMZ, A549 rTMZ) established by stepwise exposure showed no response to TMZ 10 µM alone, but NaB at 0.5 and 2.5 mM inhibited their proliferation concentration-dependently. The ratio of combination to TMZ alone was not lower in the resistant sublines than in the parental lines (A172 0.70 vs 0.86; A549 0.49 vs 0.58), indicating that the effect in resistant cells is attributable to NaB itself rather than to restored TMZ sensitivity. Millimolar exposure is the main barrier to translation.

## Introduction

Brain tumours account for a small fraction of cancer incidence in Japan but a disproportionate share of cancer death. In the national cancer registry, 5,714 brain and central nervous system tumours were recorded in 2020 against 945,055 cancers overall, yet the five-year survival of brain tumours was 35.6% against 64.1% for all cancers [1]. Glioma, including glioblastoma, accounts for roughly 4,000– 5,000 of these cases a year [2], and the five-year survival of glioblastoma itself has been reported at about 6.3% [3], placing it alongside pancreatic cancer among the most lethal malignancies.

Standard care for newly diagnosed glioblastoma has been unchanged since the 2005 randomised trial that established it [4]: maximal safe resection, followed by 60 Gy of fractionated radiotherapy with concomitant and adjuvant temozolomide (TMZ). TMZ is a triazene alkylating agent that decomposes non-enzymatically to MTIC and methylates the O6 position of guanine; the resulting mispairing drives futile mismatch repair and apoptosis. Its low molecular weight (194 Da), lipophilicity, neutral non-ionised form at physiological pH, poor recognition by efflux transporters and non-enzymatic activation together give it the central nervous system exposure that dacarbazine lacks [5, 6].

The benefit of TMZ depends on O6-methylguanine-DNA methyltransferase (MGMT). Patients whose tumours carry a methylated, and therefore silenced, MGMT promoter derive substantially more benefit from TMZ than those who do not [7]. MGMT silencing is epigenetic rather than mutational, which makes the enzymes that write and erase chromatin marks plausible drug targets in this disease [8]. Four HDAC inhibitors are in clinical use (vorinostat, romidepsin, panobinostat, chidamide), all for haematological indications.

Sodium butyrate (NaB) is a short-chain fatty acid generated by bacterial fermentation of dietary fibre in the colon and an inhibitor of class I and IIa HDACs [9]. HDAC inhibition by NaB increases histone acetylation, opens chromatin, and has been reported to induce differentiation, apoptosis and anti-inflammatory programmes across several systems: p53-independent apoptosis in HT29 colorectal cancer cells, neuroprotection through autophagy in rodent brain injury models [10], restoration of regulatory T cell function in inflammation [11], and improved insulin sensitivity in db/db mice [12]. NaB also alters splicing: RNA-seq of bovine epithelial cells treated with NaB detected 2,834 exon skipping events [13], consistent with HDAC inhibition accelerating transcription elongation.

We therefore examined NaB in glioblastoma cells, taking EGFR as a readout be-cause of its long first intron and because earlier unpublished work in this laboratory had found that SF3B inhibitors induce EGFR intron retention in A549 cells. We report the effect of NaB on EGFR expression and on proliferation in A172 glioblastoma and A549 lung adenocarcinoma cells, its combination with TMZ, and its activity in TMZ-resistant sublines derived from both lines.

## Materials and Methods

### Reagents and antibodies

Sodium butyrate (Nacalai Tesque, Kyoto, Japan) was dissolved in sterile water. Temozolomide (MedChemExpress, Monmouth Junction, NJ, USA) was dissolved in DMSO. Antibodies were anti-EGFR (4267T, Cell Signaling Technology), anti-ACTB (A00702, CGCS) and HRP-linked anti-mouse IgG (7076S, Cell Signaling Technology).

### Cell culture

A172 glioblastoma cells and A549 lung adenocarcinoma cells, the latter wild-type for EGFR, were maintained at 37 °C under 5% CO_2_ in DMEM (Nacalai Tesque) supplemented with 10% fetal bovine serum (Japan Bio Serum), MEM nonessential amino acids (×100, Nacalai Tesque) and penicillin–streptomycin (Nacalai Tesque).

### Establishment of temozolomide-resistant sublines

A172 and A549 cells were exposed to stepwise increasing TMZ concentrations. Cells were cultured for one week each at 1, 2 and 3 µM; adherent survivors were collected, diluted to one cell per well in 96-well plates, and expanded under 3 µM TMZ. The cycle was repeated at 7 and 10 µM, and the resulting clones were maintained under 10 µM TMZ. The sublines are designated A172 rTMZ and A549 rTMZ.

### Cell viability assay

Cells were seeded in 96-well plates at 2 × 10^4^ cells per well and treated with vehicle (sterile water for NaB, DMSO for TMZ), NaB, TMZ or both for up to 72 h. At 0, 24, 48 and 72 h, cells were stained using a Crystal Violet Assay Kit (Abcam) and absorbance was read at 595 nm on a microplate reader (Tecan). Values are expressed relative to the absorbance of the same condition at 0 h.

#### RT-PCR

Total RNA was extracted with TRIzol (Invitrogen) and chloroform, and 1 µg was reverse-transcribed with ReverTra Ace (Toyobo). PCR was performed with Ex Taq (TaKaRa). Primers were EGFR forward 5′-TCAGCCTCCAGAGGATGTTC-3′ and reverse 5′-GTGTTGAGGGCAATGAGGAC-3′; ACTB forward 5′-TCACCCACACTGTGCCCATCT 3′ and reverse 5′-CAGCGGAACCGCTCATTGCCAATGG-3′. Products were resolved on agarose gels.

### Western blotting

Protein was recovered by acetone precipitation from the interphase remaining after TRIzol/chloroform extraction, dissolved in urea-containing sample buffer (10 mM Tris pH 7.5, 25 mM KCl, 5 mM MgCl_2_, 6 M urea) and sonicated to shear co-precipitated nucleic acid. Protein was quantified with the Pierce 660 nm assay (Thermo Fisher). Approximately 20 µg per lane was separated by SDS-PAGE, transferred to PVDF membrane by tank blotting, blocked in Blocking One (Nacalai Tesque) for 10 min at 4 °C, incubated with primary antibody (1:1,000) for 16 h at 4 °C and secondary antibody (1:5,000) for 1 h at room temperature, and detected with Chemi-Lumi One L (Nacalai Tesque).

### Statistics

Each viability experiment was performed once. In the concentration-response experiment of Fig. 3 each condition was a single well and no dispersion is shown. In the combination experiments of Figs. 4 and 5 each condition was run in three replicate wells on one plate, and values are given as mean ± SD across those wells; differences were assessed by Student’s t-test, two-tailed and assuming equal variance, with P < 0.05 taken as significant. The groups compared are stated in each figure legend. Because the replicates are wells within a single experiment rather than independent experiments, these P values describe variation within that experiment and are reported as descriptive.

**Figure 1.**
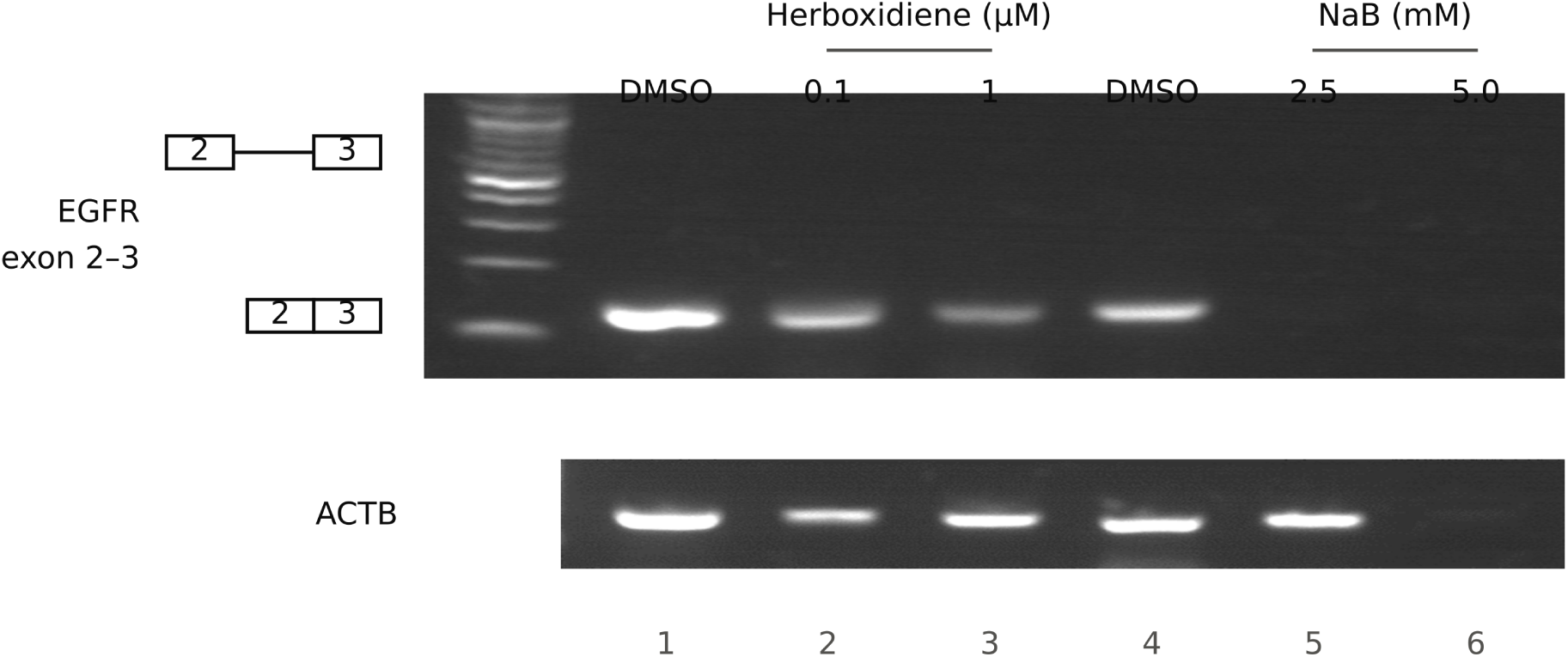
Sodium butyrate reduces EGFR expression in A172 glioblastoma cells. RT-PCR of EGFR exon 2–3 and ACTB in A172 cells treated with herboxidiene (0.1, 1 µM) or NaB (2.5, 5.0 mM). Herboxidiene modestly reduced the EGFR exon 2–3 signal without a shift in product size. NaB abolished the EGFR signal at 2.5 mM; at 5 mM the ACTB signal was also lost, indicating cell damage. Boxed exon pairs at the left mark the positions expected for the intron-retained and spliced products. Lanes are numbered below the ACTB panel.

**Figure 2.**
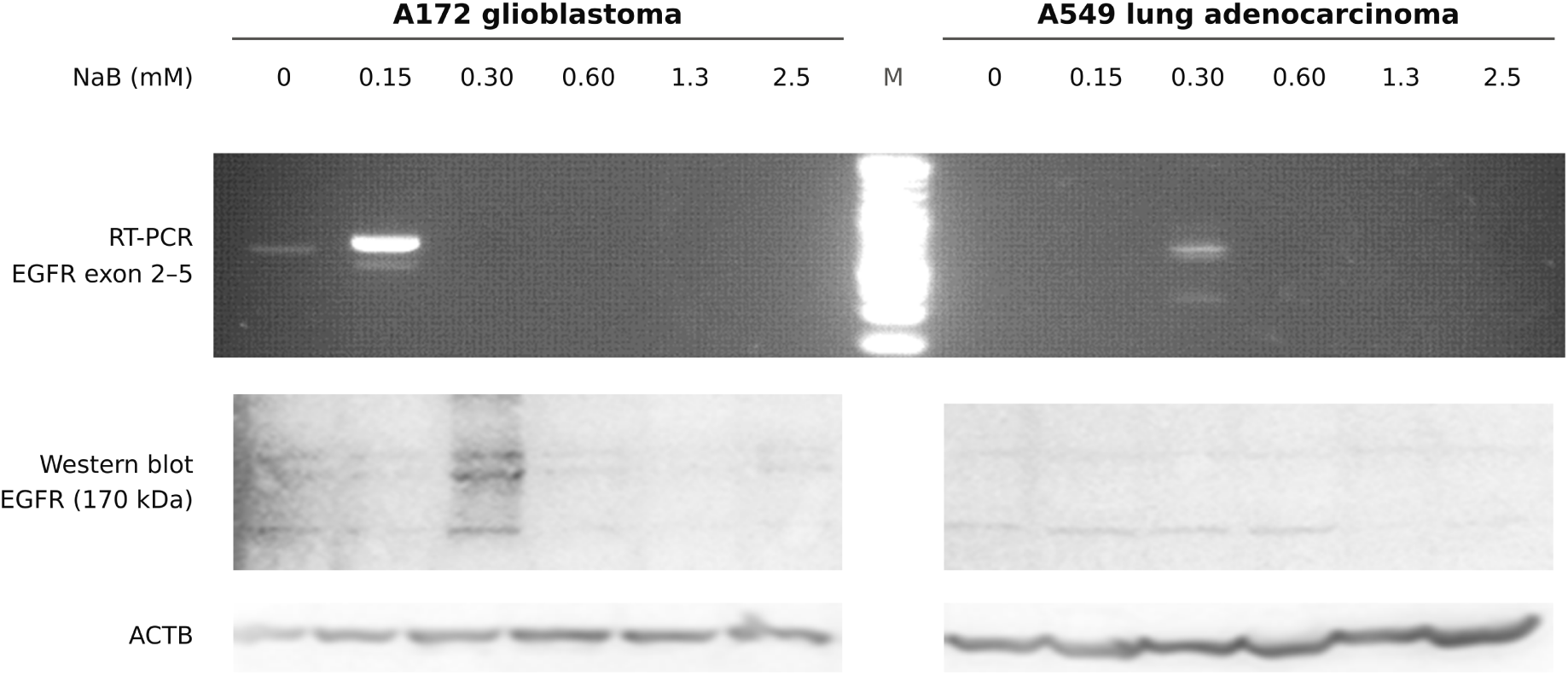
Sodium butyrate regulates EGFR expression biphasically in A172 and A549 cells. RT-PCR of EGFR exon 2–5 and western blot of EGFR (170 kDa) and ACTB after 72 h of NaB at 0, 0.15, 0.30, 0.60, 1.3 and 2.5 mM. In A172, EGFR exon 2–5 mRNA was induced at 0.15 mM and suppressed at 0.30 mM and above; EGFR protein was induced at 0.30 mM and suppressed from 0.60 mM. In A549, mRNA was induced at 0.30 mM and suppressed from 0.60 mM, with little change in EGFR protein. M, DNA size marker. The western and ACTB panels for each cell line are separate blots aligned to the lanes of the RT-PCR panel above them.

**Figure 3.**
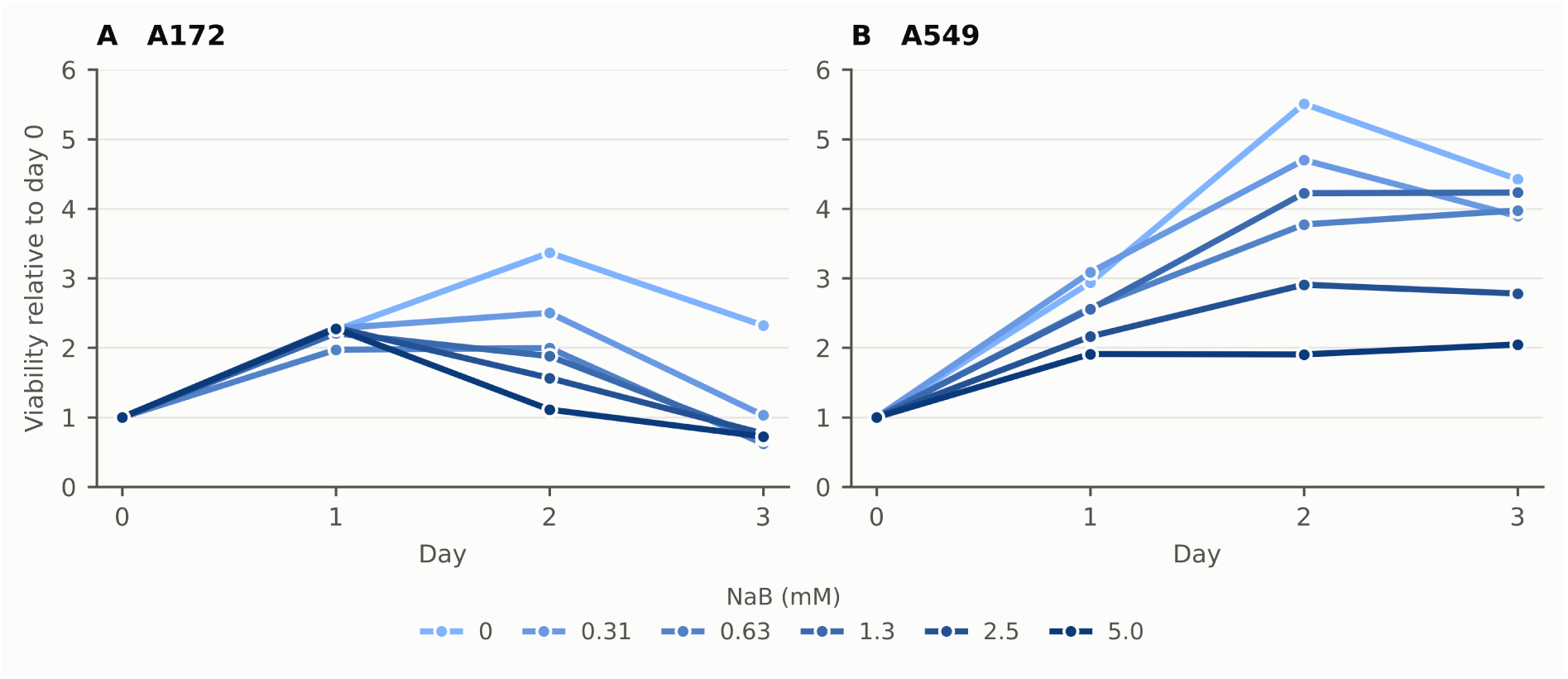
Sodium butyrate inhibits proliferation of A172 and A549 cells from 0.31 mM. Crystal violet viability assay over three days with NaB at 0, 0.31, 0.63, 1.3, 2.5 and 5.0 mM, expressed relative to day 0. (A) A172 and (B) A549 both showed concentration-dependent inhibition from day 2 at 0.31 mM and above; inhibition was weaker in A549. One experiment, one well per condition; no dispersion is shown and no statistical comparison was made. Source data are given in the supplementary file Fig3_source_data.csv.

**Figure 4.**
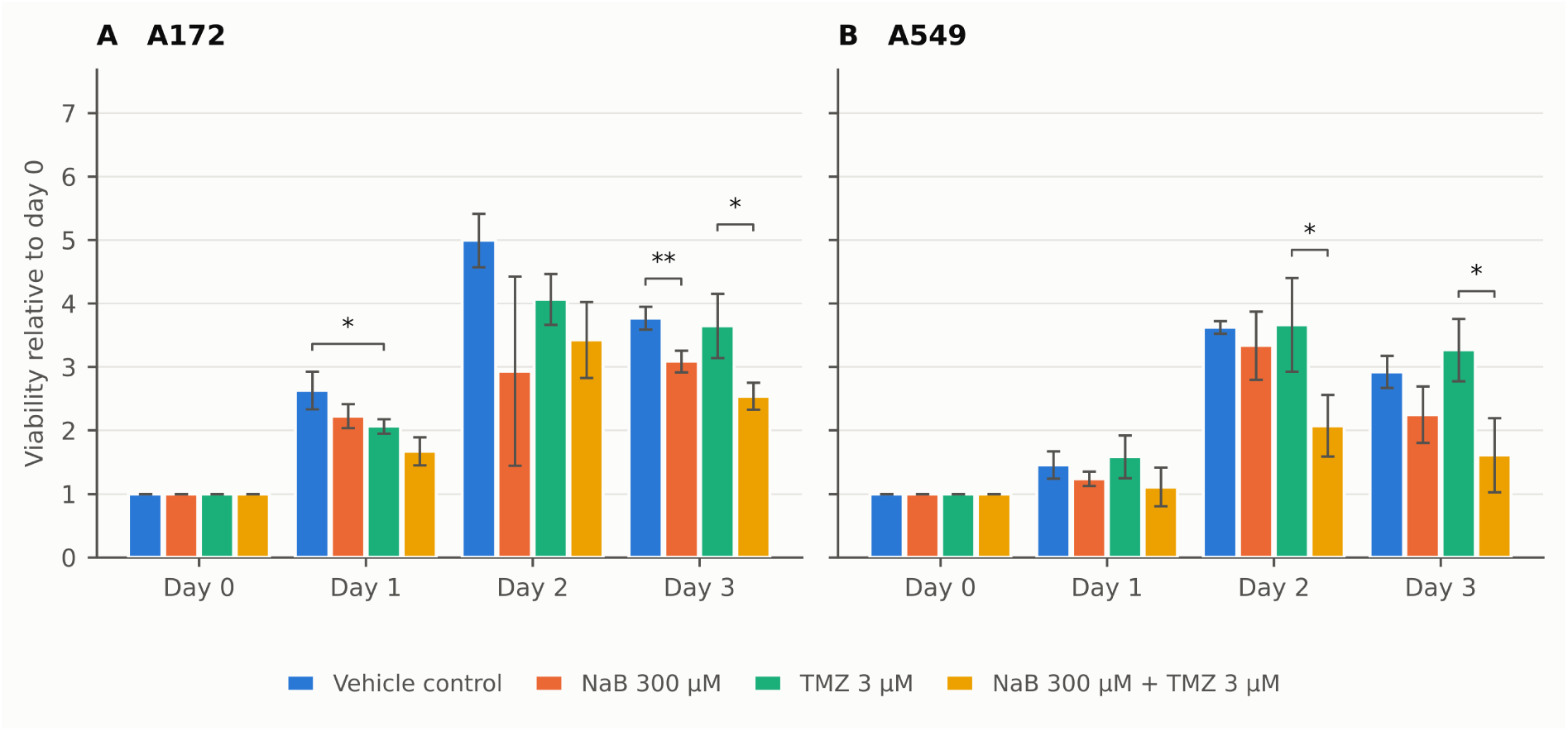
Sodium butyrate (300 µM) and temozolomide (3 µM) inhibit proliferation in combination. Crystal violet viability assay over three days with four conditions: vehicle control, NaB 300 µM, TMZ 3 µM, and both. (A) In A172, TMZ alone differed from vehicle control on day 1, NaB alone differed from control on day 3, and the combination differed from TMZ alone on day 3. (B) In A549, neither agent alone differed from control, while the combination differed from TMZ alone on days 2 and 3. One experiment; bars are the mean ± SD of three replicate wells on one plate, with the two panels on a shared y-axis. Brackets mark the two groups compared: each single agent against vehicle control, and the combination against TMZ alone. *P < 0.05, **P < 0.01 by Student’s t-test across wells. The wide SD for NaB alone in A172 on day 2 comes from one well reading 1.22 against 3.77 and 3.82; it is retained. Source data are given in the supplementary file Fig4_source_data.csv.

**Figure 5.**
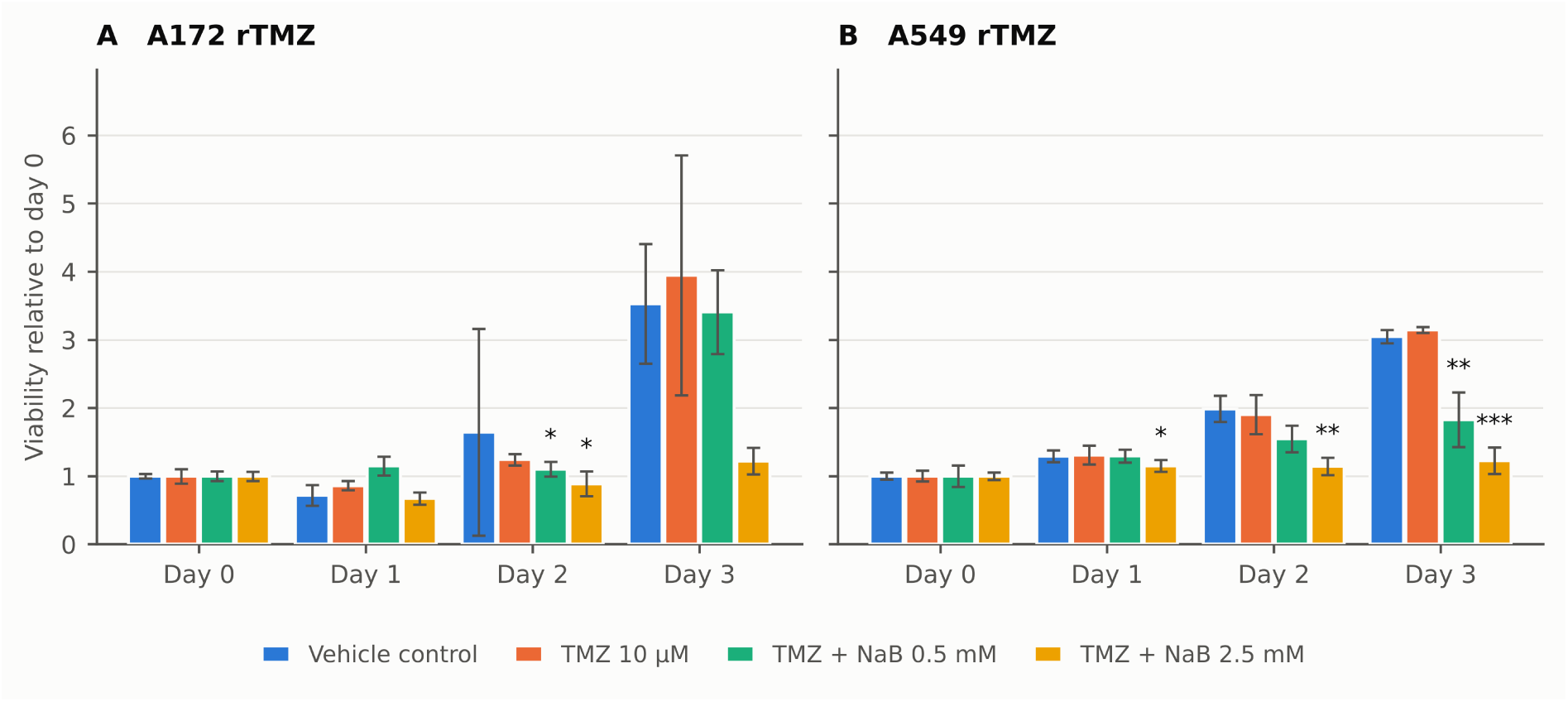
Sodium butyrate inhibits proliferation of temozolomide-resistant sublines. A172 rTMZ and A549 rTMZ cells were treated with vehicle, TMZ 10 µM alone, or TMZ 10 µM combined with NaB at 0.5 or 2.5 mM. (A) In A172 rTMZ, viability under TMZ alone matched vehicle; adding NaB lowered it on day 2 at both concentrations, while on day 3 neither comparison reached P < 0.05. The wide control error bar on day 2 comes from one well reading 3.39 against 0.84 and 0.70; it is retained. (B) In A549 rTMZ, viability under TMZ alone matched vehicle; 2.5 mM NaB lowered it on days 1 and 2, and both concentrations lowered it on day 3. One experiment; bars are the mean ± SD of three replicate wells on one plate, with the two panels on a shared y-axis. Asterisks mark the comparison against TMZ alone: *P < 0.05, **P < 0.01, ***P < 0.001 by Student’s t-test across wells. Source data are given in the supplementary file Fig5_source_data.csv.

## Results

### Sodium butyrate reduces EGFR expression in A172 cells

Earlier unpublished work in this laboratory found that the SF3B inhibitors pladienolide B and herboxidiene induce retention of EGFR intron 2 in A549 cells. In A172 cells no comparable splicing change was detected (Fig. 1). Herboxidiene reduced the EGFR exon 2–3 amplicon modestly relative to ACTB (ratio 1.71 for DMSO, 1.66 at 0.1 µM, 0.58 at 1 µM). NaB abolished the EGFR exon 2–3 signal at 2.5 mM, and at 5 mM the ACTB signal was lost as well, indicating cell damage that was not apparent on inspection of the monolayer.

### Sodium butyrate regulates EGFR expression biphasically

Because the suppression in Fig. 1 was seen only at millimolar concentrations, we examined a finer concentration series (0, 0.15, 0.30, 0.60, 1.3 and 2.5 mM) in both lines (Fig. 2). In A172, EGFR exon 2–5 mRNA was induced at 0.15 mM and suppressed at 0.30 mM and above; EGFR protein was induced at 0.30 mM and suppressed from 0.60 mM. In A549 the mRNA pattern was similar but shifted one step higher, with induction at 0.30 mM and suppression from 0.60 mM, while EGFR protein changed little. Millimolar activity is consistent with published work in prostate cancer cells, where 1 and 5 mM NaB inhibited proliferation and induced apoptosis in DU-145 and LNCaP [14].

### Sodium butyrate inhibits proliferation of both lines from 0.31 mM

Cells were treated with 0, 0.31, 0.63, 1.3, 2.5 and 5.0 mM NaB for three days and viability was measured by crystal violet staining (Fig. 3). Both lines grew rapidly, tended to reach confluence by day 2 with some detachment, and declined by day 3. From day 2 onwards NaB reduced proliferation in a concentration-dependent manner at 0.31 mM and above in both lines, more markedly in A172 than in A549. Each condition in this experiment was a single well, so the trend is descriptive and was not tested statistically. On this basis 0.30 mM was chosen for the combination experiments.

### Sodium butyrate and temozolomide inhibit proliferation in combination at individually inactive concentrations

The reported IC_50_ of TMZ in A172 is 13.7 ± 1.2 µM [15]. We therefore combined TMZ at 3 µM with NaB at 0.30 mM, concentrations chosen to produce little inhibition alone, and compared four conditions: vehicle control, each agent alone, and both (Fig. 4). In A172, TMZ alone differed from vehicle control on day 1 (P = 0.036) but not on day 3 (P = 0.72), NaB alone differed from control on day 3 (P = 0.009), and the combination differed from TMZ alone on day 3 (P = 0.025). In A549, neither agent alone differed from control at any time point, while the combination differed from TMZ alone on days 2 and 3 (P = 0.036 and 0.020).

### Sodium butyrate remains active in temozolomide-resistant sublines

A172 rTMZ and A549 rTMZ cells grew at the same rate under TMZ 10 µM as under vehicle, confirming resistance. In A172 rTMZ, adding NaB lowered viability relative to TMZ alone on day 2 at both 0.5 and 2.5 mM (P = 0.016 and 0.047); on day 3 neither comparison reached P < 0.05 (P = 0.40 and 0.058), although the mean at 2.5 mM was 1.22 against 3.95 for TMZ alone (Fig. 5A). In A549 rTMZ, 2.5 mM differed from TMZ alone on day 1 and day 2 (P = 0.025 and 0.007), and both 0.5 and 2.5 mM differed on day 3 (P = 0.003 and < 0.001), concentration-dependently (Fig. 5B). One vehicle-control well on day 2 in A172 rTMZ read 3.39 against 0.84 and 0.70 for the other two; the value is retained, and it accounts for the wide control error bar at that time point.

To ask whether resistance alters the size of the combined effect, we compared the day 3/day 0 ratio for TMZ alone with that for TMZ plus NaB (Tables 1 and 2). The combination-to-TMZ ratio was 0.70 in parental A172 and 0.86 in A172 rTMZ, and 0.49 in parental A549 and 0.58 in A549 rTMZ. The combination was therefore no more effective in the resistant sublines than in the parental lines, and the inhibition observed in resistant cells is attributable to NaB rather than to restored TMZ sensitivity. The comparison is not strictly controlled, since the two settings used different NaB concentrations (300 µM for parental lines, 500 µM for resistant sublines) and different TMZ concentrations (3 µM and 10 µM).

**Table 1:** Effect of TMZ and NaB in A172 cells by TMZ sensitivity.

| Cell line | TMZ alone (day 3/day 0) | TMZ + NaB (day 3/day 0) | Ratio (TMZ+NaB)/TMZ |
| --- | --- | --- | --- |
| A172 (NaB $300\text{ }\mu\text{M}$) | $3.65 \pm 0.51$ | $2.54 \pm 0.21$ | $0.70$ |
| A172 rTMZ (NaB $500\text{ }\mu\text{M}$) | $3.95 \pm 1.76$ | $3.41 \pm 0.62$ | $0.86$ |

**Table 2:** Effect of TMZ and NaB in A549 cells by TMZ sensitivity.

| Cell line | TMZ alone (day 3/day 0) | TMZ + NaB (day 3/day 0) | Ratio (TMZ+NaB)/TMZ |
| --- | --- | --- | --- |
| A549 (NaB 300 μM) | $3.27 \pm 0.49$ | $1.61 \pm 0.59$ | 0.49 |
| A549 rTMZ (NaB 500 μM) | $3.14 \pm 0.04$ | $1.83 \pm 0.40$ | 0.58 |

## Discussion

The clinically useful observation here is that NaB inhibits the proliferation of TMZ-resistant glioblastoma cells at concentrations at which TMZ itself has no effect. Because the ratio of combination to TMZ alone was not smaller in the resistant sublines than in the parental lines, we do not interpret this as resensitisation to TMZ; the activity belongs to NaB. For a disease in which TMZ failure leaves almost no systemic option, an agent that is indifferent to TMZ resistance is worth pursuing on its own terms.

The biphasic effect on EGFR was unexpected. Induction of EGFR exon 2–5 mRNA and protein at 0.15–0.30 mM, followed by loss of both at higher concentrations, suggests two processes with different concentration thresholds rather than a single monotonic response. HDAC inhibition opens chromatin and would be expected to increase transcription of a subset of genes; at higher concentrations, non-specific toxicity and the loss of the ACTB signal at 5 mM indicate that suppression is at least partly a consequence of cell damage. A practical implication is that the window around 0.15–0.30 mM, in which EGFR is induced, is the window in which an EGFR tyrosine kinase inhibitor might be expected to gain, rather than lose, a target. That prediction is testable in the same cell system and is the obvious next experiment.

The concentrations that inhibit proliferation are the principal obstacle. Millimolar NaB is not achievable in brain by oral dosing. Published human exposure is limited: in irritable bowel syndrome, 150 mg of microencapsulated sodium butyrate twice daily reduced abdominal pain [16]. Two routes narrow the gap. Intrathecal administration has been reported once, for leptomeningeal metastasis [17], and local delivery into the resection cavity would in principle allow tumour concentrations far above plasma. The reported half-life of butyrate is short, on the order of under an hour, so either repeated administration or a sustained-release formulation would be required; an intracranial model is needed to establish whether a tumour concentration in the required range is attainable at all.

Several limitations should be stated. Each viability experiment was performed once, so the error bars and the accompanying t-tests describe variation between replicate wells within a single experiment and not between independent experiments; the effects reported here require replication before they can be treated as established. The crystal violet assay measures adherent biomass rather than cell number, and both lines approached confluence by day 2, so the day 3 values are affected by detachment as well as by drug effect. The EGFR experiments are gel-based and semi-quantitative. Normal cells, including astrocytes and other non-transformed lines, were not tested, and the selectivity of NaB in this concentration range is therefore unknown; that comparison should precede any animal work. Finally, the combination was evaluated at single fixed concentrations of each agent, which permits a statement about the combined effect but not a formal assessment of additivity or synergy; a concentration matrix with Bliss or Loewe analysis would be needed for that.

## Supporting information

Source data for Figure 3

Source data for Figure 5

Source data for Figure 4

## Declarations

### Data availability

All data supporting the findings are contained in the figures and tables. Source data for Figs. 3, 4 and 5 are provided as supplementary CSV files. Raw images and viability data are available from the corresponding author on request.

## Acknowledgements

The authors thank the members of the Department of Clinical Pharmacology, Faculty of Pharmaceutical Sciences, Fukuoka University.

